# Developmental Trajectories of Dynamic Brain Network Organization and Their Alteration in Tourette Syndrome

**DOI:** 10.64898/2026.08.10.743841

**Authors:** Matthias Schwarz, Julia Schmidgen, Theresa V. Heinen, Azamat Yeldesbay, Nils Rosjat, Felix J. Schmitt, Kerstin Konrad, Silvia Daun, Stephan Bender

**Affiliations:** University of Cologne, Univ. Hosp Cologne, Department of Child and Adolescent Psychiatry, Germany; Forschungszentrum Jülich GmbH (INM-10), 52428 Jülich, Germany; Section Child Neuropsychology, Department of Child and Adolescent Psychiatry, Psychosomatics and Psychotherapy, University Hospital, RWTH Aachen, Germany; Institute of Medical Psychology and Medical Sociology, Faculty of Medicine, RWTH Aachen University, 52074 Aachen, Germany; Cognitive Neuroscience, Institute of Neuroscience and Medicine (INM-3), Research Centre Jülich, 52248 Jülich, Germany; Computational Systems Neuroscience, Institute of Zoology, University of Cologne, Germany; Computational Neuroscience – Modeling Neural Network Function, Institute of Zoology, University of Cologne, Germany

**Keywords:** Developmental trajectories, Tourette syndrome, source connectivity states, theta band, network flexibility, tic severity, source connectivity, network maturation

## Abstract

Typical brain network maturation involves an increase in network flexibility and hemispheric specialization. Tourette syndrome (TS) disrupts these trajectories, with tic severity potentially modulating deviations. This study examined theta-band EEG source connectivity states in typically developing children and children with TS. We assessed age-related trajectories and the impact of tic severity using generalized linear modeling, accounting for sex and multiple comparisons. K-means clustering identified four recurrent source connectivity states (A–D), with metrics including Coverage, representing state prevalence (proportion of time spent in each state), Average Dwell Time, an index of state stability (mean duration of stable persistence of each state), and Transition Rate Per Minute, reflecting global network flexibility (frequency of state switches per minute). In healthy controls (HC), typical maturation was characterized by increased left intra-hemisphere connectivity state stability and prevalence, decreased diffuse connectivity state stability, and rising network flexibility. TS patients exhibited deviant trajectories, including age-dependent decreasing global network flexibility across subgroups stratified by tic severity and marginally divergent diffuse activity patterns, with high-severity cases showing increased diffuse connectivity state stability. The normative patterns suggest typical motor development requiring dynamic network reconfiguration and hemispheric specialization, processes that appear altered in TS. TS patients exhibit age-dependent network rigidity across severity subgroups, as reflected by decreased transition rates, alongside severity- modulated network imbalances, indicating that tic disorders disrupt mechanisms of brain network maturation underlying motor control. These findings suggest that atypical trajectories of network stability and flexibility represent a key feature of tic pathophysiology, highlighting the role of altered network dynamics in TS during maturation.

## 1 Introduction

Childhood and adolescence represent a critical period in neurodevelopment and motor maturation, characterized by large-scale reorganization in functional network architecture (Denckla, 1974; Haywood & Getchell, 2021; López-Vicente et al., 2021). Developmental studies have demonstrated that during this developmental window, motor skills undergo significant changes driven by maturational processes that enhance coordination, speed, and precision of motor actions (Denckla, 1974; Rueda et al., 2004; Schmidgen et al., 2024). While normative developmental trajectories can vary between individuals, neurodevelopmental disorders are often associated with measurable alterations in network maturation (Duan et al., 2021a; Nishida et al., 2013).

Such deviations from normative motor network development were particularly observed in disorders like Gilles de la Tourette syndrome (TS) (Franzkowiak et al., 2012; Schmidgen et al., 2025). TS is classified as a tic disorder, which is among the most common neuropsychiatric disorders during childhood and adolescence, and is characterized by sudden, involuntary, non-rhythmic motor or vocal tics (Cohen et al., 2013; Knight et al., 2012; Müller-Vahl, 2019). For all subclassifications of tic disorders, lifetime prevalence is estimated to be between 1 and 3%, while purely motor tics are more common than vocal or combined tic disorders like TS (Cohen et al., 2013; Knight et al., 2012; Spencer et al., 1995). Symptom onset is typically at ages around 6 to 8 years, while maximum symptom severity is typically around 10 to 12 years, with a 60% to 80% symptom remission in adolescence (Black et al., 2016; Ricketts et al., 2022).

Symptom severity in TS can differ greatly between individuals, impairing social interactions as well as general daily functioning. Children with high tic severity experience significantly more limitations of self-efficacy as well as academic performance, which are linked to higher psychological impairments (Bloch et al., 2006; Cravedi et al., 2017; Eapen et al., 2016). Studies have shown a correlation between tic severity and a higher risk of comorbid psychiatric disorders like attention deficit hyperactivity disorder, obsessive-compulsive disorder, and affective disorders, impeding psychological and social adaptation (El Malhany et al., 2015; Kumar et al., 2016; Leckman et al., 2014). In social contexts, tic disorders are often misinterpreted as intentional misbehavior, which can contribute to the stigmatization and isolation of patients (Eapen et al., 2016). The Yale Global Tic Severity Scale (YALE) has been established as a reliable tool for standardized assessment of tic severity and is clinically validated for initial assessment and longitudinal evaluation of frequency, intensity, complexity, and functional impairment of vocal and motor tics (Leckman et al., 1989, 2014; Storch et al., 2005). Specifically, the YALE motor-subscore (YALE-m) was validated and showed high reliability in reflecting motor tic severity (Storch et al., 2005). Recent studies confirm that YALE not only accurately reflects overall tic severity but also demonstrates robust responsiveness to clinical change in intervention studies (Leckman et al., 1989, 2014). Importantly, children and adolescents with higher YALE scores are more likely to experience school-related difficulties, peer rejection, and reduced quality of life compared to those with milder symptoms (Eapen et al., 2016; Ricketts et al., 2022).

At this point, the underlying mechanisms of TS are not yet fully understood, hindering progress in developing therapeutic measures (Albin, 2018; Knight et al., 2012; Yael et al., 2015). Previous studies suggest that, from a pathophysiological perspective, TS is linked to a disbalance within cortico-striato-thalamo-cortical circuits, where impaired motor neuron inhibition leads to the emergence of tics (Franzkowiak et al., 2012; Heise et al., 2010; Houghton et al., 2014; Mink, 2003; Rae & Critchley, 2022; Worbe et al., 2015). Furthermore, compensational patterns in prefrontal regions indicate the development of adaptive tic suppression mechanisms (G. M. Jackson et al., 2015; S. R. Jackson et al., 2011; Mueller et al., 2006; Rae & Critchley, 2022). Another hypothesis is based on findings that individuals with TS tend to experience premonitory urges and hypersensitivity, indicating deficits in the integration of sensory input and motor output. This concept describes an alternated perception-action binding, where subjects with TS show an enhanced motor response to sensory input (Beste et al., 2016; Friedrich et al., 2021; Kleimaker et al., 2020; Petruo et al., 2020).

A key aspect of network maturation involves theta band oscillations (4-8 Hz), which encode essential motor development processes and sensory-motor integration (Beste et al., 2023; Cruikshank et al., 2012; Tomassini et al., 2017). The association between theta band activity and motor development has been demonstrated in previous studies (Ahn et al., 2022; Schmidgen et al., 2025; Wendiggensen et al., 2023). Specifically, resting-state EEG applications have demonstrated the ability to reveal insights into dynamic networks by profiling functional connectivity over time. This method offers key advantages for studying pediatric brain network dynamics, including task-free assessments that minimize performance and motivation confounds, making them especially suitable for children and adolescents (De Pasquale et al., 2010; Fox & Raichle, 2007; Fries, 2015; Tarailis et al., 2024). By calculating microstates from continuous resting-state EEG recordings, dynamic brain configurations can be characterized (Beckmann et al., 2005; Koenig et al., 2024; Seitzman et al., 2017; Tarailis et al., 2024). Longitudinal investigations demonstrate high test reliability of resting-state microstate features, which represent a dynamic view of the spatial distribution of the electric potential on the scalp over time, as well as dynamic measures of functional connectivity. This emphasizes their potential as reproducible biomarkers in neurodevelopmental research (Hommelsen et al., 2022; Khanna et al., 2014). Disruptions in specific microstates for neuropsychiatric disorders, such as dementia, depression, schizophrenia, and TS were found (Nishida et al., 2013; Stevens et al., 2007; Strik et al., 1995; Wang et al., 2021). While conventional EEG microstate frameworks analyze brain activity on sensor level, Rosjat et. al. (2024) introduced a novel framework allowing functional connectivity mapping on the cortical level using source connectivity states. For functional connectivity state measurements, a precise quantification of functional coupling between brain regions in resting-state EEG can be calculated using the corrected imaginary Phase-Locking Value (ciPLV), which eliminates volume conduction artifacts and identifies true phase coupling between source time series (Brunia et al., 2011; Rosjat et al., 2024; Tenke & Kayser, 2012). From this measurement, temporarily stable connectivity states reflecting network structures can be extracted through cluster analyses and then used to study motor development based on their progression characteristics (Rosjat et al., 2018, 2024). Building on this framework, the present study utilizes EEG connectivity state analysis to provide further insights into the changes and developments in motor networks shown in these developmental studies (Rosjat et al., 2018, 2024).

Previous studies have examined either the age-related development of EEG-based network dynamics or the group differences between tic patients and control subjects (Duan et al., 2021a; Morand-Beaulieu et al., 2025; Rosjat et al., 2024), yet integrative investigations that simultaneously consider normal development and pathological deviations are rare. In particular, it remains unclear whether and how resting-state connectivity metrics evolve with age, and whether these developmental trajectories diverge between tic patients and healthy controls or vary by tic severity. To address this research gap, this study investigates theta band connectivity during resting-state eyes-closed conditions. It aims to: (1) characterize ciPLV-based connectivity states in motor development across childhood and adolescence; (2) profile normative patterns of motor maturation in order to identify potential deviations from regular development; (3) compare these dynamic metrics between tic patients and controls; and (4) examine variation within the patient cohort as a function of tic severity. Considering previous findings on network maturation during motor development in theta-band oscillation and their alterations in TS, this study specified the following hypotheses:

1. Theta resting-state ciPLV connectivity state analysis will show systematic age-related changes during motor development in children and adolescents, following a developmental trajectory with age.
2. Children and adolescents with TS will exhibit divergent age-related developmental trajectories in theta resting-state ciPLV connectivity state analysis.
3. Within the patient cohort, these age-dependent developmental trajectories will be modulated by tic severity, as measured by the YALE motor-subscore (YALE-m).

Overall, this study aims to strengthen the bridge between neurophysiological research and clinical symptoms, ultimately optimizing the understanding of pathophysiology in children and adolescents with tic disorders.

## 2 Methods

### 2.1 Participants

This study includes data of 28 patients with a diagnosed TS (female n=7, age (mean ± sd): 10.58 ± 2.41 years, range: 7.40 - 16.6 years) and 52 healthy control (HC) subjects (age (mean ± sd): 11.01 ± 3.19 years, range: 5.25 - 16.6 years). Specific to the TS cohort, each subject had a preexisting TS diagnosis by DSM-5. We confirmed the given DSM-5 criteria of TS in the patient group and conducted the K-DIPS interview with every participant and their parent or guardian to check the absence of comorbidities except attention deficit and hyperactivity disorder (ADHD) in the patient group, as well as all psychiatric disorders in the HC cohort (Fleischhaker et al., 2011; Margraf et al., 2017). In both groups, participants were excluded (i) if a full-scale IQ below or equal to 70 was evaluated by the WISC-V (Wechsler, 2024), (ii) if born before 32 weeks gestation, (iii) if they or a parent had a history of epilepsy or other central nervous system disorders, (iv) if they had uncorrectable visual impairments, (v) if they had taken psychoactive substances before the assessments (24 hours for caffeine and nicotine, 48 hours for stimulants, four half-lives for tranquilizers or antipsychotics), (vi) if they had metallic or electrical implants, (vii) if they experienced severe back pain, extreme obesity, or were pregnant. As this study primarily focuses on motor cortical development, tic severity was evaluated with the clinician-administered and verified motor tic-specific YALE-m, which rates motor tics along with their associated functional impairment (Leckman et al., 1989; Storch et al., 2005).

### 2.1 Experimental setup

Electrophysiological data were acquired using a BrainAmp 64-channel DC amplifier (Brain Products GmbH, Munich, Germany) and BrainProducts Easycaps with Ag/AgCl sintered electrodes in an equidistant layout. Impedances were kept below 5 kΩ, and all channels were referenced to Cz. Eye movement artifacts were monitored using four electrooculogram (EOG) electrodes (two horizontal electrooculogram channels placed at the outer eye canthi (HEOG1, HEOG2) and two vertical electrooculogram channels placed infra-orbital (I01, I02). Signals were digitized at a sampling rate of 5000 Hz using Brain Vision Recorder (Brain Products GmbH, Munich, Germany). Structural MRI data were collected at the Research Centre Jülich using a 3-Tesla Siemens MAGNETOM Prisma scanner (Siemens Healthcare, Erlangen, Germany). High-resolution T1-weighted images were obtained with a rapid gradient echo sequence, employing a repetition time of 1790 ms, an echo time of 2.53 ms, and an 8° flip angle. A total of 176 slices were acquired (slice thickness 0.9 mm, interslice gap 0.45 mm) over a 256 mm field of view, yielding isotropic voxels of 0.9 x 0.9 x 0.9 mm.

### 2.2 Experimental protocol and paradigm

After EEG setup and electrode preparation, every subject had one recording session lasting around 20 minutes. This included four consecutive 120-second resting-state blocks, alternating between eyes-closed (EC) and eyes-open (EO) conditions (two blocks each), providing comprehensive data while accommodating attention spans. During the measurements, the participants sat comfortably in a dimly lit room and were instructed to remain as still as possible and to relax. The participants were asked to close their eyes gently without falling asleep. Between each recording, a short break was included, and the research assistant verbally announced the next recording, giving the participants corresponding instructions. Electrode impedances were verified to remain below 5 kΩ throughout the session. Besides the EEG recording, an approximately six-minute-long structural MRI measurement was conducted in a separate session at the Research Center Jülich.

### 2.3 EEG preprocessing

EEG preprocessing was conducted using MATLAB, EEGLAB, and MNE-Python, as well as our own Python scripts (Delorme & Makeig, 2004; Henschel et al., 2020; Inc, 2022). Continuous data was first downsampled to 512 Hz to reduce computer utilization, while still retaining higher frequency band activity. Filtering, drift correction, artifact removal, independent component analysis, and bad channel interpolation were conducted using the RELAX Junior pipeline (Hill et al., 2024). A zero-phase bandpass filter from 0.25 to 80 Hz and a 50 Hz notch filter were applied to suppress line noise. EOG channels were used to locate muscle or eye movement artifacts and were ultimately excluded from the dataset. Subsequently, the data were re-referenced using average referencing before concatenating the data to continuous trials. Because recordings from children are more prone to artifacts when eyes are open, only EC data were analyzed to maintain maximal comparability. Finally, each dataset underwent a thorough visual inspection to confirm its quality.

### 2.4 MRI preprocessing

Preprocessing of the structural MRI scans was performed using FastSurfers recon-all pipeline (Henschel et al., 2020). The raw T1-weighted scans were automatically corrected for inhomogeneities, aligned, and normalized. Afterwards, skull-stripping as well as pial surface reconstruction is conducted to receive surfaces of gray matter, white matter, inner skull, outer skull, and outer skin. Finally, a three-conductivity (scalp =0.33 S/m, skull ≥0.0042 S/m, brain ≥0.33 S/m) boundary element model (BEM) was constructed for each subject using MNE-Python. Referencing predetermined anatomical landmarks by using own Python scripts, each electrode montage was individually adjusted to the corresponding BEM model to ensure precise source localization. After each step, the MRI data were visually inspected using FastSurfers integrated visualization tool. Misaligned montages were manually corrected.

### 2.5 Source localization

For each subject, the source space was constructed using FastSurfer’s BEM model and the individually adjusted electrode montage. An octant-6 source spacing ensured at least 3.5mm distance to the skull surface. After performing forward modelling using MNE-Python, source localizations were categorized in 62 parcels according to the Desikan-Killiany-Tourville Atlas (Klein & Tourville, 2012). Within each parcel, the predominant orientation was identified, and opposite-pointing vertices were inverted so that all source estimates aligned consistently. Afterwards, time courses of all vertices of a parcel were averaged to obtain a unified parcel time series. This procedure was harmonized through close cooperation with studies from the Research Centre Jülich, where it was verified by EEG recordings of different task paradigms (Rosjat et al., 2024).

### 2.6 Phase-locking connectivity analysis

After source localization, our approach to construct time-resolved functional networks is segmented into (a) connectivity analysis and (b) graph construction, closely aligned with the framework of the Dynamic Synchronization Toolbox (Rosjat & Daun, 2022). The canonical frequency band Theta (4-8 Hz) was used to isolate neurophysiologically meaningful oscillatory components.

Each parcel’s time course was loaded as a two-dimensional NumPy array, with one dimension corresponding to parcels and the other to time points. EEG signals were band-pass filtered in the theta range (4-8 Hz) using a zero-phase finite-impulse-response (FIR) filter (Hamming window design, filter order automatically determined by MNE). This approach allowed only the frequency of interest to remain while preserving the phase of the signal. The analytic signal of each filtered time series was then computed via the Hilbert transform, and its argument yielded the instantaneous phase ϕ(t) at every time point t.

To capture the rapid transitions that characterize EEG source connectivity states, the corrected imaginary Phase-Locking Value (ciPLV) was computed using a 300 ms sliding-window approach with 200 ms overlap, which produces an effective time resolution of 100 ms between successive windows (Rosjat et al., 2024). This approach matches the intrinsic timescale of EEG microstates,which have been shown to remain quasi-stable for about 100 ms before shifting to a new pattern (Koenig et al., 2024; Lehmann et al., o. J.; Tarailis et al., 2024)

ciPLV was chosen for its ability to quantify phase synchronization while largely excluding zero-lag contributions that may arise from volume conduction or common reference effects, making it a widespread standard in studies of resting-state networks and task-evoked dynamics, where the identification of reliable, non-zero-lag synchronization is critical for mapping directed information flow and network topology (Brodbeck et al., 2012; Bruña et al., 2018a; Rosjat et al., 2024). Traditional measures such as the Phase-Locking Value (PLV) capture the overall consistency of phase differences but cannot distinguish genuine interactions from spurious coupling at zero phase lag. In contrast, ciPLV isolates the imaginary part of the complex phase-difference distribution, effectively discounting instantaneous (zero-lag) synchronization that is likely artifactual. By focusing on the imaginary component, ciPLV retains sensitivity to true time-lagged interactions, reducing the risk of spatial leakage, signal mixing, and thus misleadingly high coupling estimates at zero phase difference (Bruña et al., 2018b). Formally, for two parcels m and n with instantaneous phases *φ_m_*(*t*) and *φ_n_*(*t*) over *T* time points, ciPLV is defined as:

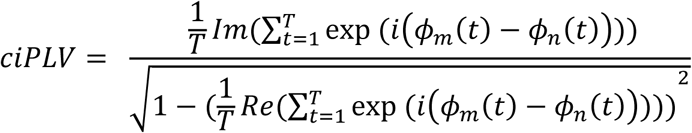

To make the resulting time courses comparable, we normalized upon obtaining the window-wise ciPLV matrices. We first computed the mean absolute ciPLV over all windows for each parcel pair and then subtracted this mean from every window’s absolute value. This centers each pair’s time course around zero, so that only deviations from its overall coupling strength remain. Diagonal entries were set to zero to eliminate self-coupling artifacts and prevent them from dominating subsequent analyses. A small constant ε = 1x10^-10^ was used to avoid division by zero.

To distill significant network structure from the dense coupling data, we used the Brain Connectivity Toolbox to retain only the top 10% of strongest connections in each window (Rubinov & Sporns, 2010). This sparsification emphasizes robust interactions, reduces noise, and improves interpretability, in line with prior recommendations for functional brain networks (Rubinov & Sporns, 2010; Sporns, 2013). Connections above or equal to the 90th percentile threshold were assigned a value of one, while all others were set to zero, producing a binary adjacency matrix for each time point. This way, each matrix defines an undirected graph *G_t_* = (*V*, *E_t_*), where *V* = {1, … , *P*} is the set of parcels and *E_t_* the set of edges that define the strongest connections. By concatenating the graphs *G*_1_, … , *G_W_* we constructed a dynamic network representation that captures evolving patterns of synchrony at a resolution of 100ms. An overview of the anaylsis steps is illustrated in Figure 2.

**Fig. 1.**
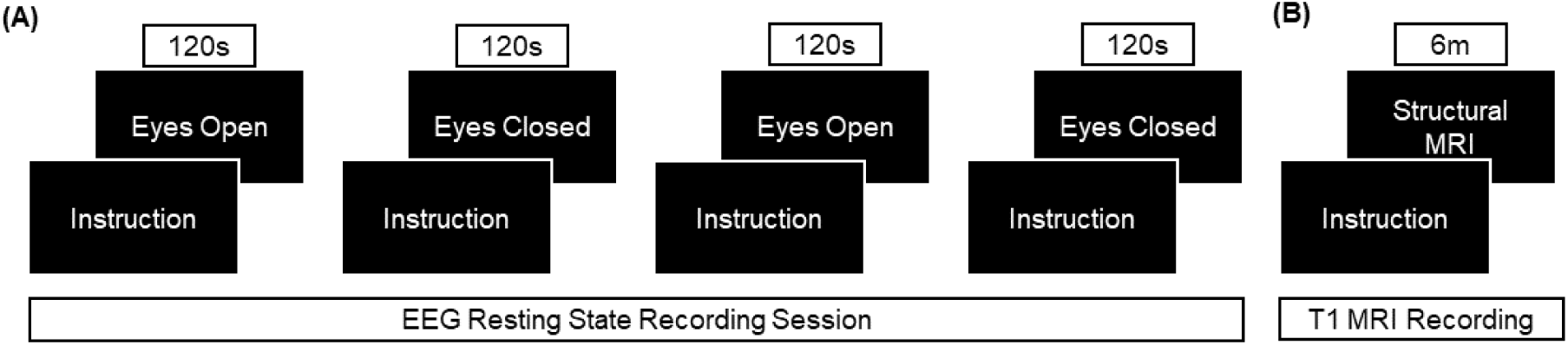
Experimental design. **(A)** A single measurement run consisted of four 120s resting-state EEG recordings alternating in eyes-open and eyes-closed condition. **(B)** On a separate visit, participants underwent a structural T1-weighted MRI scan lasting about 6 minutes. *Note*. EO = Eyes Open; EC = Eyes Closed; EEG = Electroencephalogramm; MRI = Magnetic Resonance Imaging

**Fig. 2.**
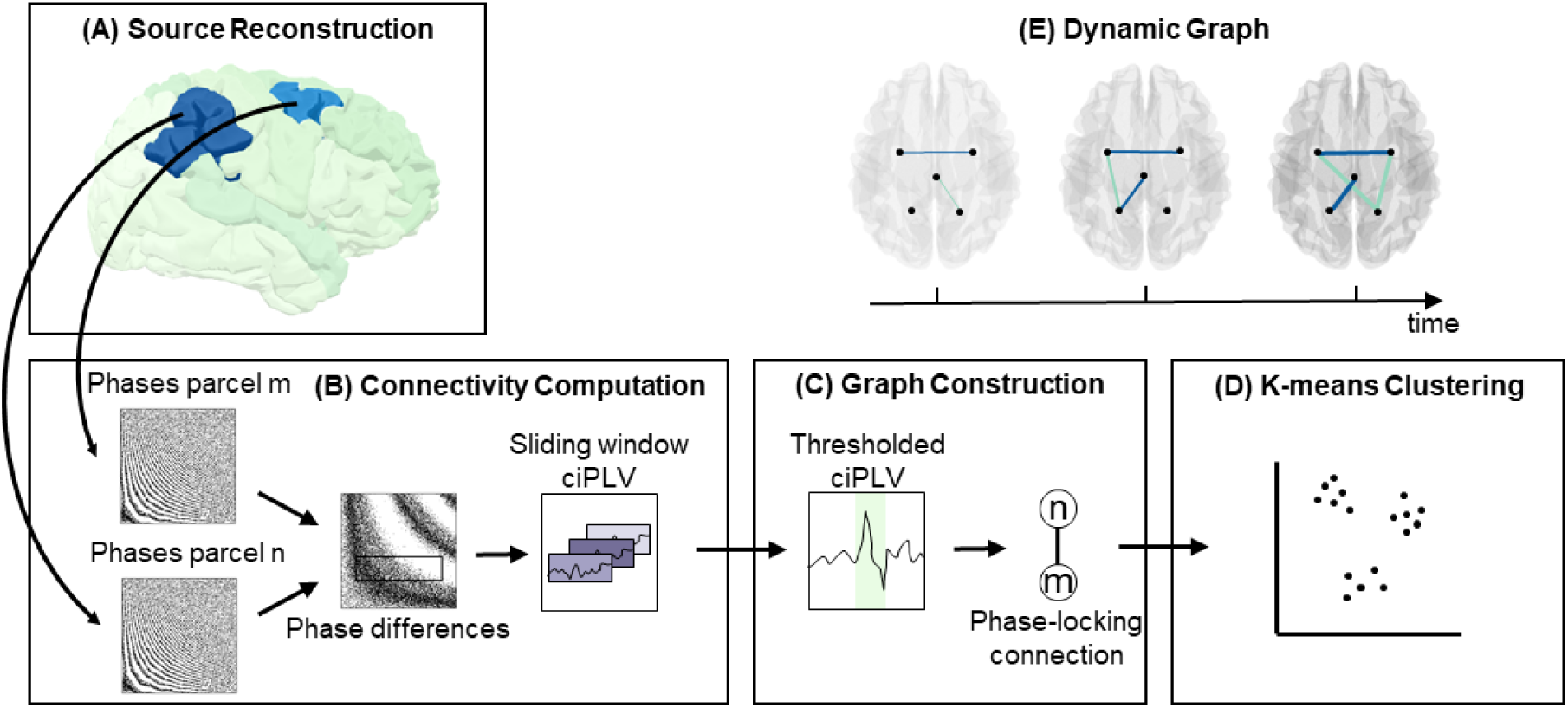
Illustration of the procedure for calculating connectivity states, adapted from Rosjat et al. (2024), Fig. 2. **(A)** For any pair of regions (m and n), source activity is extracted, and **(B)** ciPLV is computed using theta-band phase differences within timeframes of 100 ms. **(C)** For each window, ciPLV is thresholded at 90 %, retaining only the strongest connections between regions. **(D)** Resulting connectivity patterns are clustered across all region pairs and time frames using K-means, **(E)** creating a dynamic source-based graph representation.

### 2.7 Connectivity state clustering

To cluster the connectivity patterns into discrete connectivity states, we applied K-means clustering to the concatenated ciPLV time-window matrices derived from eyes closed resting state trials across the theta frequency band. For analyses of healthy development, clustering was performed on the dataset of 52 HC, whereas for subsequent comparative analyses, K-means clustering was applied to a combined dataset comprising 28 patient recordings and 52 HC sessions. Before clustering, we employed the elbow criterion to assess the optimal number of clusters, which supported a four-state solution, substantiating our choice and aligning our approach with previous studies (Rosjat et al., 2018, 2024).

Sample weights were assigned inversely proportional to the number of time windows contributed by each participant, thereby equalizing the influence of each subject in the overall clustering solution. Window counts were tallied using NumPy’s unique function, and weights were computed as the reciprocal of these counts. We configured the K-means estimator with four clusters, twenty restarts, and a fixed random seed to guarantee reproducibility. After fitting the model on the z-transformed feature matrix, we extracted cluster labels for every time window and retrieved the standardized centroids, which were then back-transformed into the original connectivity scale for subsequent interpretation.

By mapping each time window to one of the four identified connectivity states, we were able to compute a suite of temporal metrics at the individual level, as described in previous studies used to quantify both microstates and source connectivity states (Rosjat et al., 2024; Britz et al., 2010; Koenig et al., 1999). These metrics included:

1. Coverage: The relative proportion of time spent in each state, calculated by the number of windows in the state divided by the total number of analyzed windows, representing state prevalence;
2. Average Dwell Time: The mean duration a state remains stable before a transition into another state occurs, calculated by the sum of time spent in a state divided by its occurrences, an index of state stability;
3. Transition Rate Per Minute: The total number of transitions in the recorded session, normalized per minute of recording, reflecting global network flexibility.

Finally, the four cluster centroids were visualized as circular connectivity plots, where each centroid matrix was rendered with region labels arranged according to their angular positions, as well as projected onto a three-dimensional brain model.

Figure 3 shows the four source connectivity states extracted from the data. The observed spatial patterns closely resemble configurations prior reported for EEG source connectivity states (Rosjat et al., 2024). Specifically, we observed the following principal states:

a. An intrahemispheric connectivity pattern within the left hemisphere (State A, “left hemispheric”)
b. A mirrored intrahemispheric pattern within the right hemisphere (State B, “right hemispheric”)
c. A parietal-dominant bilateral connectivity configuration (State C, “parietal-dominant”)
d. A more diffuse and globally weaker connectivity pattern with a slight frontal predominance (State D, “diffuse”)

**Fig. 3.**
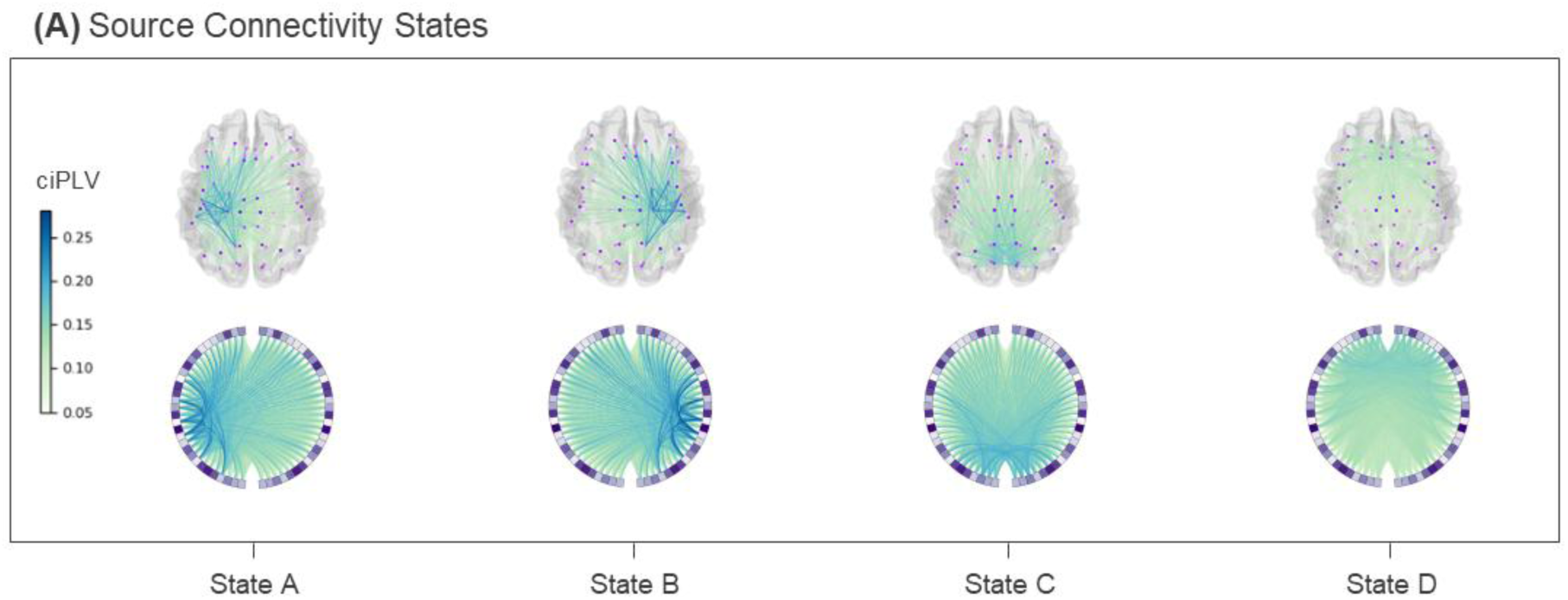
Source connectivity states computed across all participants in the EC condition **(A)**, showing (from left to right) the previously described left-hemispheric state A, right-hemispheric state B, parietal-dominant state C, and diffuse state D. *Note*. EC = eyes-closed

Finally, each time window for every participant was assigned to one of these four source connectivity states, enabling the computation of individual temporal metrics. We calculated four metrics, including Coverage (the relative proportion of windows assigned to a given state), Average Dwell Time (the mean uninterrupted duration in a state) and Transition Rate per Minute (the count of state switches relative to the recording length).

### 2.8 Statistical analysis

All statistical analyses were conducted in Python using the Statsmodels and SciPy libraries (Seabold & Perktold, 2010; Virtanen et al., 2020), with heteroskedasticity-consistent standard errors applied throughout to guard against heteroscedasticity. As our outcomes are bounded, partly right-skewed, and exhibit mean-variance coupling, we opted for Generalized Linear Models (GLMs). In order to assess these distributional properties and ensure data quality, the data were visually examined before testing. Across all analyses, the distribution of the dependent variables dictated the model family: For positive, right-skewed measures such as Average Dwell Time or Transition rate, we fitted GLMs from the Gamma family with a log link. For proportion-type outcomes, including Coverage of individual EEG states, we used Gaussian GLMs on logit-transformed data. To account for group imbalance, sensitivity analyses using a randomly selected matched subsample of the HC group were conducted, confirming the robustness of the primary results. In these models, centered age served as the primary predictor, and sex was included as a covariate. Ordinary Least Squares (OLS) models specified analogously to the GLMs later produced estimates and test statistics highly consistent with the GLM outputs.

Before model fitting, we prepared covariates to capture both linear and nonlinear developmental effects. Age was mean-centered and further transformed to create quadratic and cubic terms. However, higher-order polynomials did not improve model fit. Therefore, we specified age as a linear term to preserve interpretability and parsimony.

First, to establish normative developmental trajectories, we analyzed the control group in isolation. Subsequently, to identify deviations associated with TS, group differences between patients and controls were then examined in the full sample using a GLM that incorporated main effects of group and centered age, their interaction, and sex. Dependent variables were modeled with the same families, Gamma with a log link for continuous positive measures, and Gaussian with logit-transformed outcomes for proportions, as in the developmental analyses. This approach allowed us to test for both baseline differences and potential age-related divergences between groups.

Given the clinical heterogeneity inherent in TS, we investigated whether group-level trajectories were uniformly representative of the entire patient cohort. To account for potential masking effects where low- and high-severity patients might exhibit divergent developmental trends, we conducted a targeted post-hoc sensitivity analysis using median-split severity stratification. Based on the median YALE-m score, patients were divided into low-YALE-m and high-YALE-m subgroups. Subsequently, we fitted separate GLMs comparing each subgroup against the HC group. This stratified approach served to distinguish between global diagnostic and severity-driven alterations.

Multiple comparison correction was uniformly applied across all analyses using the Benjamini-Hochberg false discovery rate (FDR) procedure. We reported two-sided FDR-adjusted p-values as q with a threshold of q<0.05 unless otherwise noted. This analytic framework allowed us to exploratively examine normative age effects, group differences, and symptom-related modulation of dynamic EEG connectivity with appropriate attention to distributional properties and multiple testing. While all results are FDR-adjusted, they remain preliminary; thus, outcomes should be interpreted with care.

## 3 Results

To characterize the development-dependent network dynamics, the metrics Average Dwell Time and Coverage, as well as the Transition Rate across all states, were calculated for all four identified connectivity states (States A–D). Only the significant modulations are reported below.

### 3.1 Cortico-cortical development in healthy controls (HC)

We first examined age-related changes in the metrics by computing theta-band connectivity states exclusively within the HC group (n=52). The four connectivity states identified in HC were nearly identical to the ones observed using data from all participants, showing that the identified states reflect stable and reproducible patterns of functional connectivity. Sex-adjusted GLMs across ages 5-16 years revealed that multiple metrics tracked cortico-cortical maturation in HC (see Figure 4).

**Fig. 4.**
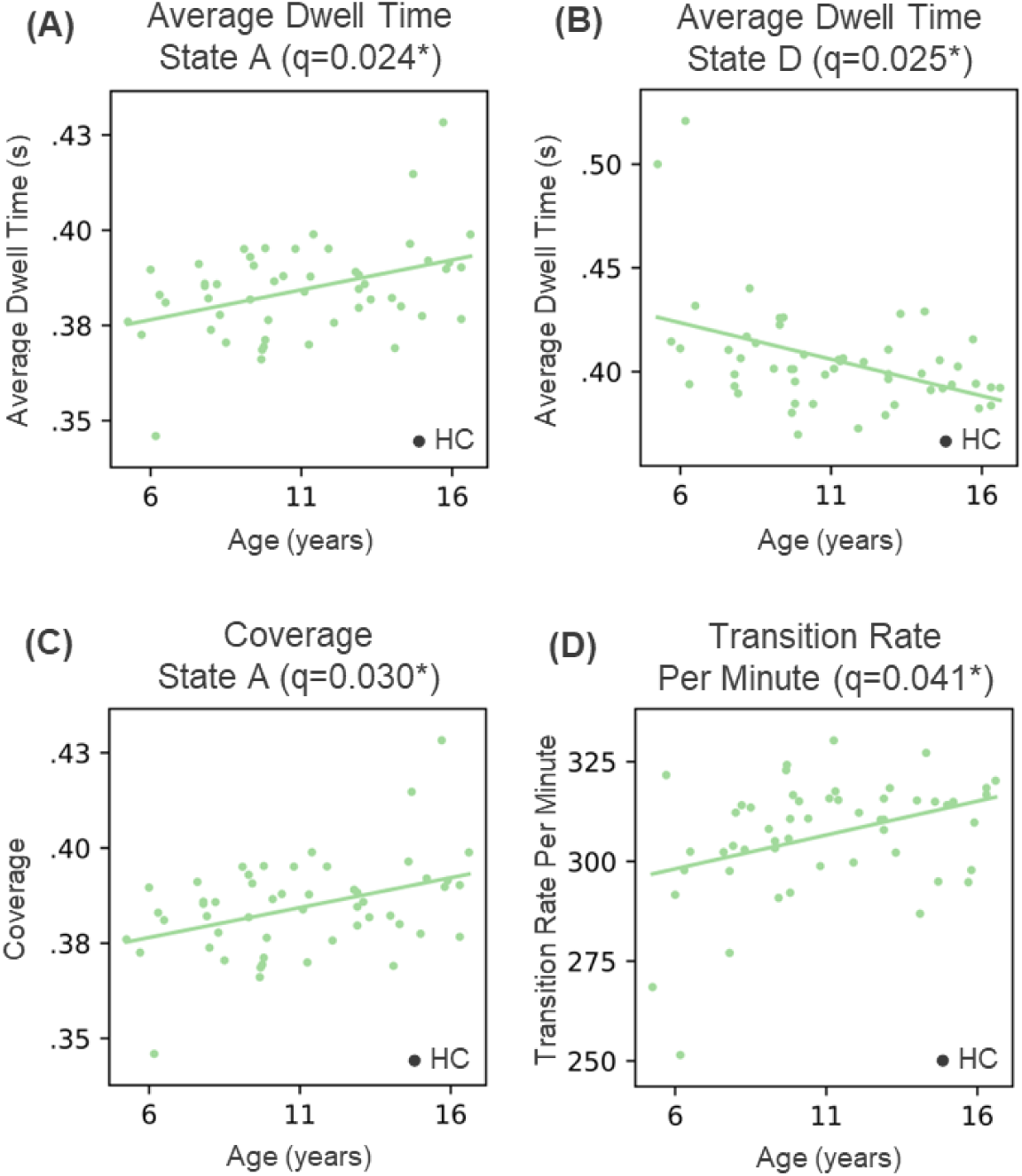
Developmental trajectories of theta oscillation during EC resting state in HC. **(A)** Regression plots representing GLM analyses of age-dependence in Average Dwell Time of left-hemispheric state A, **(B)** diffuse state D, as well as **(C)** Coverage of left-hemispheric state A, indicating a physiological increase in network stability by usage of higher connectivity networks and decrease in diffuse connectivity. **(D)** Regression plot showing the result of a GLM analysis of Transition Rate Per Minute in the EC condition, suggesting higher network flexibility with age in HC. HC = healthy controls; EC = eyes-closed; GLM = General Linear Model

Several state metrics showed significant maturation in the HC group: In particular, the average duration of time spent in the left-hemispheric state A increased with age (Fig. 4(A), RR/year = 1.004, 95 % CI [1.001, 1.007], q = 0.024). A convergent age-related increase was observed in the coverage of the same state (Fig. 4(C), RR/year = 1.031, 95 % CI [0.007, 0.0054], q = 0.041). These findings show increased left-hemispheric state A prevalence and stability in childhood maturation of HC. In contrast, the diffuse state D decreased in mean duration with increasing age (Fig. 4(B), RR/year = 0.992, 95 % CI [-0.015, -0.002], q = 0.025). Transition Rate Per Minute was observed to increase with age for HC (Fig. 4(D), RR/year = 1.006, 95 % Cl [0.001, 0.010], q = 0.030). In summary, these findings indicate physiological maturation characterized by a stabilization of localized networks with simultaneously increased flexible switching ability.

### 3.2 Between-group differences in EEG resting-state metrics

To investigate developmental differences in EC resting-state theta oscillations among children and adolescents with TS, an analysis of theta band connectivity states was conducted using the full EEG dataset. Sex-adjusted GLMs were then applied to previously described metrics in order to examine group differences in state stability, usage, and overall network flexibility. Two specific patterns emerged, which can be interpreted as global and severity-associated markers, respectively.

In contrast to the age-dependent increase in Transition Rate observed in HC, the TS group exhibited a significant divergent trajectory (RR/year = 0.990, 95% Cl [-0.017, -0.004], q = 0.019). Specifically, TS patients showed a stagnation or reduction in network flexibility with advancing age, as visualized in Fig. 5(A). The post-hoc median split analysis confirmed that this effect was consistently present in both the low-YALE-m (RR/year = 0.987, 95 % CI [- 0.020, -0.005], q = 0.016) and the high-YALE-m subgroup (RR/year = 0.980, 95 % CI [-0.013, -0.002], q = 0.021) (Fig. 5(B)). These findings suggest that reduced maturational increase in network flexibility is a fundamental and severity-independent characteristic of the TS phenotype.

**Fig. 5.**
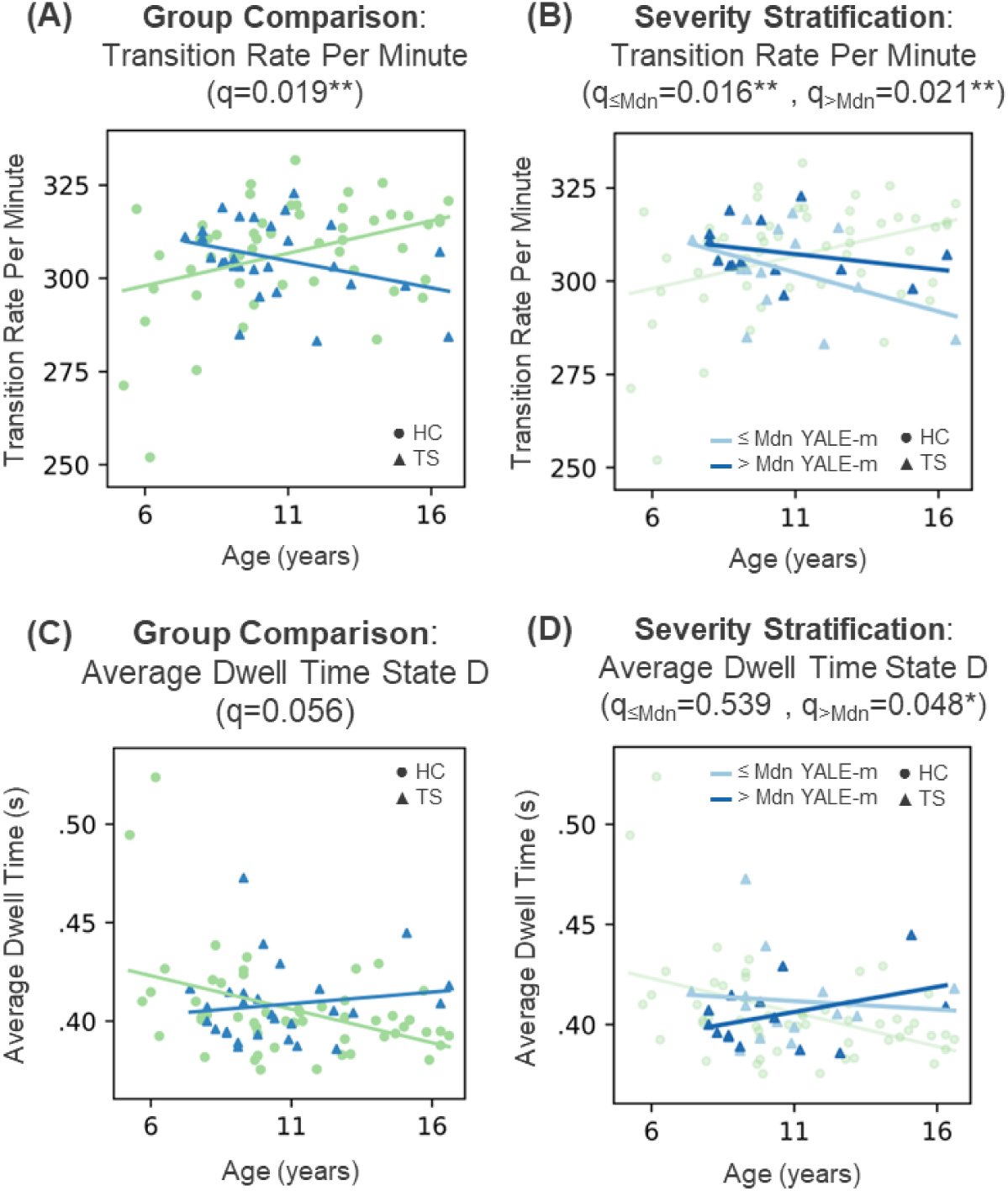
Between-group differences in theta connectivity during EC resting state. **(A)** Regression plots of the GLM analyses visualizing deviances of developmental trajectories for Transition Rate Per Minute: While HC exhibit an age-dependent increase in network flexibility, TS patients show a decline. **(B)** Median split stratification by tic severity (YALE-m) showing reduced Transition Rate Per Minute across both low- and high-severity subgroups, suggesting this metric as a possible diagnostic marker. **(C)** GLM analysis of Average Dwell Time in diffuse state D shows a trend towards divergent trajectories, where diffuse activity stagnates or rises in TS compared to the physiological decrease in HC. **(D)** Stratification in low- and high-severity subgroups reveals a significant deviation only in TS with high tic severity, identifying the persistence of diffuse activity as a potential severity-driven marker. Mdn = Median, YALE-m = YALE Global Tic Severity Scale motor score, HC = healthy controls, TS = Tourettes syndrome, EC = eyes-closed, GLM = General Linear Model

Analysis of diffuse state D revealed a contrasting age-related trajectory in Average Dwell Time between groups (RR/year = 1.010, 95 % CI [0.001, 0.018], q = 0.056). Specifically, Average Dwell Time decreased with age in HC, but remained stable or showed a slight increase with age in patients (Fig. 5(C)). While group mean comparison only showed a statistical trend, median split analysis differentiated this picture: Whereas the low-YALE-m patients showed no significant difference compared to the HC group, only high-YALE-m patients showed a significant deviation in this developmental pattern (RR/year = 1.012, 95 % CI [0.002, 0.023], q = 0.048) (Fig. 5(D)). The results suggest that the persistence of diffuse activity patterns is associated with higher tic severity.

## 4 Discussion

### 4.1 Summary of findings

In this study, we examined resting-state theta band EEG source connectivity states. K-means clustering revealed four recurrent source connectivity states (Fig. 3), each showing distinct age-related trajectories and tic-severity effects. In the HC group (5-16 years), linear age analyses revealed typical developmental trajectories, such as an increase in average duration of time spent in the left-hemispheric state as well as an increase in state coverage for the same state. Mean duration of diffuse activity decreased with age in HC, whereas network flexibility, measured by the rate of state transitions, increased during maturation. In contrast to this normative increase of network flexibility, between-group comparisons revealed that children and adolescents with TS exhibited a significant age-dependent decrease of state transition rate across both the low- and high-severity subgroups. Furthermore, while in HC, the mean duration of diffuse activity decreased with age, group comparison indicated a marginally divergent trajectory in TS. Median-split sensitivity analysis revealed a significant deviation from normative development only in the high-severity subgroup, in which the Average Dwell Time of the diffuse state increased with age.

### 4.1 Normative maturation of theta dynamics

The observed increase of transitions per minute over age, thus higher network flexibility, is consistent with developmental models that predict networks to mature from local, short-range configurations to more flexible and integrated configurations (Fair et al., 2009, Hill et al., 2023), accompanied by greater top-down connectivity (Hwang et al., 2010). This suggests enhanced inter-network exchange with increasing age in healthy development. An age-related increase in the left-hemispheric state coverage with age, as well as a decline of the diffuse state, aligns with findings of reductions in salience- and attention-related network involvement, as well as an age-dependent decrease of weaker connections (Bagdasarov et al., 2022; Deery et al., 2023), indicating a transition toward more stable large-scale networks with increasing age. Overall, our findings reinforce the prevailing view that normative brain maturation involves a shift away from weakly integrated, non-modular states and toward a predominance of well-modulated network states.

### 4.2 Reduced network flexibility in TS

The age-related increase in transition rates in healthy controls suggests that normative brain maturation is associated with enhanced network flexibility and network integration (Hill et al., 2023). Our findings revealed that children with TS did not exhibit this normative increase in state transition rates, but showed stable or decreasing rates, indicating diminished network flexibility and slower reconfiguration capacity. This atypical developmental trajectory is consistent with reports of immature functional brain organization in TS (Church et al., 2009; Openneer et al., 2020). Converging evidence from task-based EEG studies further supports this interpretation: Prochnow et al. (2025) reported enhanced theta-band modulation during perception–action binding tasks in TS, indicative of stronger and more rigid coupling between sensory input and motor output. In line with our resting-state findings of reduced state transition rates, heightened perception–action binding, and immature functional organization, suggests overly rigid coupling between functional systems (Schmidgen et al., 2025; Kleimaker et al., 2020). Such inflexible network alignments may constrain the brain’s ability to dynamically disengage and reconfigure functional states, thereby facilitating repetitive and stereotyped motor output.

Notably, this deviation from normative brain maturation was observed in both low- and high-severity TS, suggesting that reduced network flexibility reflects a trait-like characteristic of TS rather than a marker of symptom severity. Accordingly, network flexibility, as indexed by transition rates per minute, may represent a global neurodevelopmental feature of TS and a potential diagnostic or early risk marker.

### 4.3 Maturational lag in diffuse state activity as a marker of tic-severity

While the network flexibility deficit was severity-independent in TS patients, the stability of diffuse activity observed in state D followed a dose-dependent pattern. Our findings regarding healthy development align with studies supporting that healthy maturation, low-integration patterns are pruned in favor of specialized networks (López-Vicente et al., 2021). The results of this study suggest that while low-severity TS patients largely follow this normative developmental trajectory, patients with higher motor tic severity exhibit a maturational lag, characterized by the persistence of diffuse activity states. This prolonged maintenance of diffuse network configurations may be driven by deficits in synaptic plasticity, which constrain synaptic pruning and delay the emergence of stable, specialized functional networks (Felling & Singer, 2011). Such a pattern could reflect a propensity for more diffuse connectivity patterns in TS, supporting previous evidence of disrupted long-range network integration in individuals with TS (Duan et al., 2021b). Importantly, impaired long-range integration may limit efficient communication between control, sensorimotor, and associative networks, thereby exacerbating difficulties in motor inhibition and top-down regulation.

### 4.4 Methodological strengths

To ensure consistency and reproducibility, the methodological framework of this study was carefully aligned with established approaches from comparable peer-reviewed studies (Rosjat et al., 2018, 2024). The inclusion of 28 tic patients and 52 healthy controls represents a comparatively large cohort, increasing the interpretability of the results. Source-level EEG using individualized BEM head models and dSPM reconstruction reduces volume-conduction effects, allowing detailed investigation of 62 cortical regions. Particularly, the use of ciPLV minimizes zero-lag artifacts (Bruña et al., 2018a). Selecting the theta band as our primary focus reflects convergent findings central to tic disorders and motor development, as demonstrated by recent neurophysiological investigations (Prochnow et al., 2025; Schmidgen et al., 2025; Takacs et al., 2024; Wendiggensen et al., 2023). The topological characteristics of the identified source connectivity states closely align with previous findings in EEG source connectivity literature (Rosjat et al., 2024). The three predefined metrics used to quantify these states are scientifically established for the quantification of microstates as well as source connectivity states (Bagdasarov et al., 2022; Rosjat et al., 2024; Rubinov & Sporns, 2010). Finally, the use of a linear age term is consistent with multiple comparable studies with childhood populations, preserving the interpretability under limited sample size conditions (Bagdasarov et al., 2022; Hill et al., 2023; Jiang et al., 2024; Kang et al., 2024).

### 4.5 Limitations

Several methodological and conceptual limitations should be acknowledged to contextualize the present results. General limitations to our study are the cross-sectional nature of the sample, restricting the interpretability of age-related associations and developmental conclusions, as well as our methodological decision to exclusively focus on the theta band, which may overlook effects in other relevant frequency bands. Furthermore, while ciPLV reduces zero-lag artifacts, special leakage remains a possible source of error. Potential confounding effects of comorbidities must be considered, given their link to reduced default-mode and frontoparietal network disruption (Lin et al., 2015; Uddin et al., 2008). However, the primary limitation of this study is the restricted sample size and age distribution of the patient cohort, with n=28 where 19 patients were aged between 8 and 11 years. This type of distribution limits the stability of interactions between age and group, as well as age and YALE-m significantly, therefore results should be carefully contextualized. A larger, more continuous, and ideally longitudinal sample is needed to confirm developmental trajectories and robustly model non-linear age trends.

### 4.6 Conclusion

Our findings demonstrated that normative network maturation during motor development in children and adolescents is characterized by higher network flexibility, increased left-hemispheric network usage, and a shift away from diffuse activity with increasing age. TS patients inherited an age-dependent reduction in network flexibility compared to typically developing peers, while symptom severity modulated an increase with diffuse state activity over age, indicating disruptions in large-scale network maturation in patients with tic disorders. To translate and consolidate these findings, future research should combine a larger and more age-balanced sample cohort with a longitudinal framework to validate the described age interactions. Furthermore, combining dynamic source-based EEG with fMRI recordings could deepen the insight into the multi-scale network signature in tic disorder. These cross-modal approaches may yield a further understanding of divergent motor development within TS.

## Acknowledgments

We acknowledge the fruitful collaboration with the CRC 1451 projects A06 and B03. Furthermore, we thank Felix J. Schmitt and the ITCC (IT Center University of Cologne) for assisting with and providing computing resources on the DFG-funded high-performance computing system RAMSES (Research Accelerator for Modeling and Simulation with Enhanced Security).

## 5 Data and Code Availability

The analysis code is available at https://github.com/matthschwa/dynamic-brain-networks-ts.git for review and adaptation. Due to ethical restrictions, the underlying data can be obtained from the corresponding author upon reasonable request. Data access is subject to ethical approval and may require formal agreements, and full reproducibility will only be possible once the data are released under these conditions.

## 6 CRediT authorship contribution statement

**Matthias Schwarz:** Conceptualization, Methodology, Visualization, Formal analysis, Writing – original draft**. Julia Schmidgen:** Investigation, Data curation, Validation, Conceptualization, Methodology, Formal analysis, Writing - Review & Editing. **Azamat Yeldesbay**: Validation, Software. **Nils Rosjat:** Software. **Theresa Valentine Heinen:** Methodology, Conceptualization. **Felix J. Schmitt:** Validation, Formal analysis. **Kerstin Konrad:** Resources, Project administration, Conceptualization, Funding acquisition. **Stephan Bender:** Funding acquisition, Supervision, Project administration, Resources.

## 9 Ethics approval statement

This study was conducted in accordance with the local legislation and institutional requirements. The research protocol was reviewed and approved by the Ethics Committee of the University of Cologne and the Ethics Committee of the University of Aachen.

## Abbreviations

ADHD: Attention deficit and hyperactivity disorder
BEM: Boundary element model
Ci: Confidence interval
ciPLV: Corrected imaginary Phase-Locking Value
DC: Direct current
DSM-5: Diagnostic and Statistical Manual of Mental Disorders, Fifth Edition
EC: Eyes-closed
EEG: Electroencephalogram
EO: Eyes-open
EOG: Electrooculogram
FDR: False discovery rate
GLM: Generalized linear model
HC: Healthy controls
Hz: Hertz
K-DIPS: Structured Diagnostic Interview for Mental Disorders in Children and Adolescents
MRI: Magnet resonance imaging
OLS: Ordinary least squares
PLV: Phase locking value
RR: Relapsed rate
SE: Standard error
TS: Tourette syndrome
WISC-V: Wechsler Intelligence Scale for Children, Fifth Edition
YALE: The Yale Global Tic Severity Scale
YALE-m: YALE motor-subscore

